# Intraductal patient derived xenografts of estrogen receptor positive (ER+) breast cancer recapitulate the histopathological spectrum and metastatic potential of human lesions

**DOI:** 10.1101/409177

**Authors:** Maryse Fiche, Valentina Scabia, Laura Battista, Assia Treboux, Athina Stravodimou, Khalil Zaman, RLS, Ayyakannu Ayyannan, George Sflomos, Cathrin Brisken

**Affiliations:** International Cancer Prevention Institute, CH-1066 Epalinges, Switzerland; Swiss Institute for Experimental Cancer Research, School of Life Sciences, Ecole Polytechnique Fédérale de Lausanne, CH-1015 Lausanne, Switzerland; Centre Hospitalier Universitaire Vaudois, University Hospital of Lausanne, CH-1011 Lausanne, Switzerland

**Keywords:** intraductal xenografts, luminal breast cancer, preclinical model, patient-derived xenografts, ductal carcinoma *in situ*, metastasis

## Abstract

Estrogen receptor α positive (ER+) or “luminal” breast cancers were notoriously difficult to establish as patient-derived xenografts (PDXs). We and others recently demonstrated that the microenvironment is critical for ER+ tumor cells; by grafting them into milk ducts >90% take rates are achieved and many features of the human disease are recapitulated. This intra-ductal (ID) approach holds promise for personalized medicine, yet human and murine stroma are organized differently and this and other species specificities may limit the value of this model. Here, we analyzed 21 ER+ ID-PDXs histopathologically. We find that ID-PDXs vary in extent and define four histopathological patterns: flat, lobular, *in situ*, and invasive, which occur in pure and combined forms. The ID-PDXs replicate earlier stages of tumor development than their clinical counterparts. Micrometastases are already detected when lesions appear *in situ*. Tumor extent, histopathological patterns, and metastatic load correlate with biological properties of their tumors of origin. Our findings add evidence to the validity of the intraductal model for *in vivo* studies of ER+ breast cancer and raise the intriguing possibility that tumor cell dissemination may occur earlier than currently thought.

**Conflict of interest statement:** The authors declare no conflict of interest.

## Introduction

Breast cancer (BC) is a frequent disease worldwide [1]. Over 75% of BCs express the estrogen receptor α (ER) in >1% of the tumor cells by immunohistochemistry (IHC) [2]. ER+ BCs largely overlap with the luminal subtypes defined by global gene expression [3,4]. Luminal BC is heterogeneous and further subdivided into luminal A and B BC, with low *versus* high proliferative indices and distant recurrence rates [5]. Twenty percent of patients experience distant recurrence and cancer-related death [6]. Over-treatment of the early stages of the disease and endocrine resistance are additional problems in this subgroup [7]. A lack of preclinical models hampered progress in understanding the biology of these tumors and the development of new therapeutic approaches. Genetically engineered mouse models mostly develop hormone receptor negative tumors, and few ER+ BC cell lines grow *in vivo*, and only if supplemented with nonphysiological amounts of estrogen [8]. PDXs are increasingly used for preclinical studies but difficult to establish from ER+ tumors [8]. We and others showed that the microenvironment is a major determinant of luminal BC cells and that take rates increase dramatically when hormone sensitive BC cells are grafted to mouse milk duct [9]. They grow in the absence of estrogen supplementation, recapitulate biological and molecular features of their clinical counterpart and respond similarly to therapy [9,10]. Yet, mammary stroma and endocrine milieu differ beween women and mice. To assess the impact of the mouse host on the biology of the engrafted human BC cells of this increasingly used model, we analyzed 21 ID-PDXs histopathologically.

## Materials and Methods

The study was approved by the Commission cantonale d’éthique de la recherche sur l’être humain (CER*-* VD 38/15), patients signed an informed consent and all animal experiments were performed in accordance with protocols approved by the Service de la Consommation et des Affaires Vétérinaires of Canton de Vaud, Switzerland. Within 30 minutes of surgical excision, after inking of margins and macroscopic assessment, a fraction of tumor tissue was taken by the pathologist (MF) and transported to the laboratory in DMEM/F12. Information on patient age, menopausal status, primary tumor (PT) size, nodal status, histological type according to WHO 2012 classification [11], ER, progesterone receptor (PgR), Ki67, HER2 status, was collected from clinical charts and anonymized (Table S1).

Tumor pieces were mechanically and enzymatically dissociated to single cells, lentivirally transduced with luciferase-GFP and injected into the teats of 10-week-old immunocompromised female mice (NSG) [9]. *In vivo* growth was monitored every two weeks by measuring bioluminescence. Engrafted glands were dissected, fixed in buffered formalin for 2 hours and paraffin-embedded. Fifteen 4 μm sections were cut and nr 1, 7, and 15 stained with Hemalun Eosin (H&E). ALU staining was used to unequivocally identify human cells when required. IHC was performed under standard clinical conditions with Discovery Ventana ULTRA [9]. Metastatic load was calculated as percentage of bioluminescence positive organs of organs collected post mortem.

## Results

Tumor cells from 21 patients were ID grafted to 88 mice in 220 glands after lentiviral transduction with luciferase-GFP (Figure 1A, B, Table S1). Mice were sacrificed when radiance >10E8. Presence of micrometastases was determined by *ex vivo* radiance measurements on distant organs revealing bones as the most frequent site of tumor cell seeding followed by lungs, brain and liver (Figure 1C). Tumor extent was evaluated semi-quantitatively on H&E stained sections of engrafted glands; it varied with ≥70% of the ducts distended by human cells in 6/21 PDXs (Figure S1A), tumor cell foci occupying 20-60% or < 5% of the ductal tree in 10 and 5 PDXs, respectively (Figure S1B, C).

**Figure 1:**
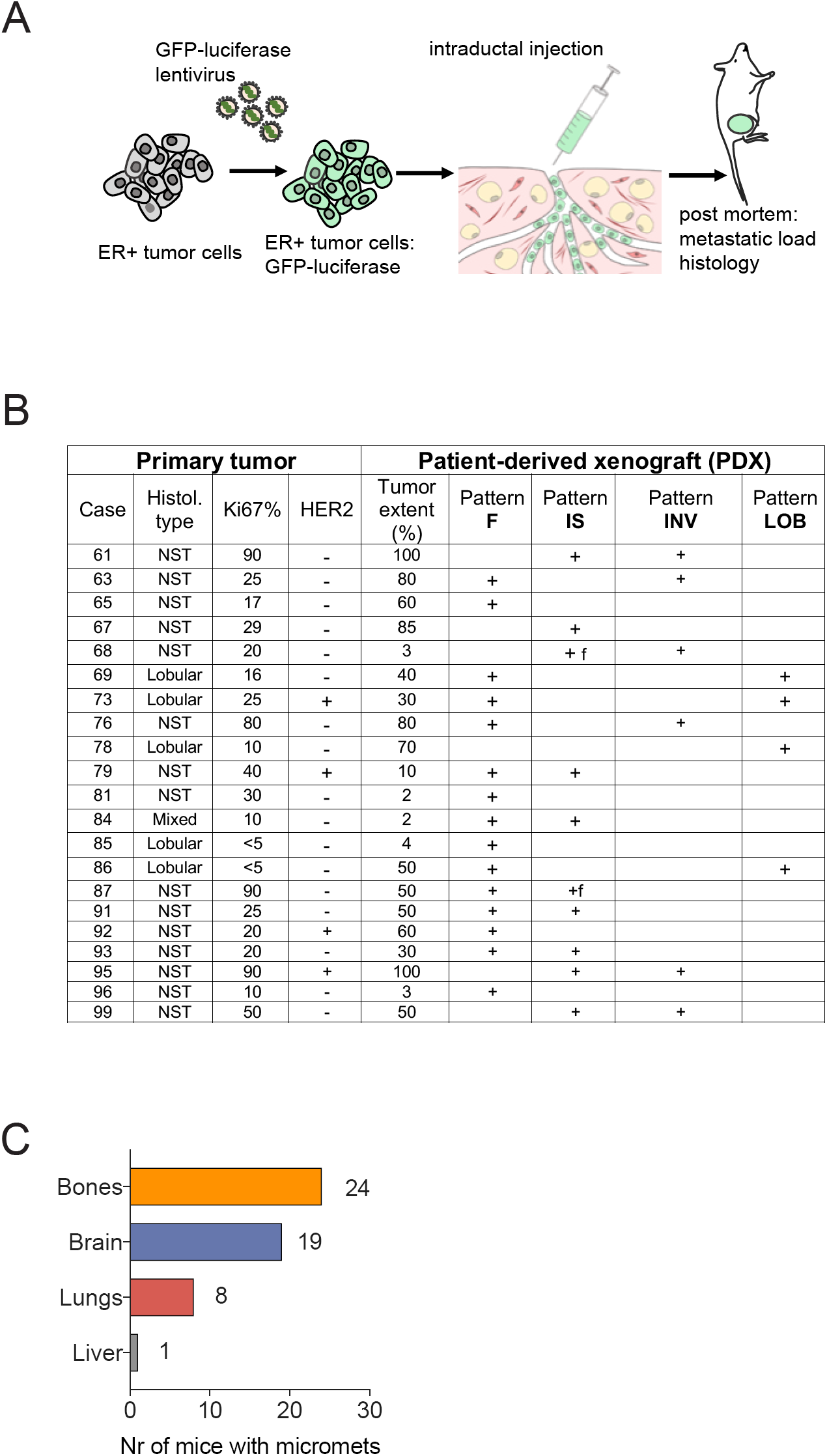
Intraductal ER+ breast cancer patient-derived xenografts. (A) Experimental scheme of PDX establishment and follow-up. PTs are dissociated to single cells, which are subsequently transduced with lentivirus encoding GFP and luciferase. The infected cells are injected intraductally in multiple glands and their *in vivo* growth is monitored by bioluminescence. The mice are euthanized and the presence of metastases is assessed by bioluminescence in distant organs. (B) Table reporting PT and PDX characteristics. F: foci. (C) Bar graph showing number of xenografted mice (n=88) bearing micrometastases in different target organs.

ER expression was detected in all 21 PDXs by IHC and associated with PgR expression in 17/21 PT (80.9%) and in 12/21 PDX (57.1%) (Table S1). Overall, the distribution of ER+ and PgR+ cells in PDXs and PTs were similar (Figure S2A, B) but in 5 and 16 pairs, respectively, positivity was discrepant (Figure S2C, D).

HER2 status was IHC 3+ in 2 of 3 PDXs corresponding IHC 3+ PTs including one confirmed by FISH. A PT with a IHC 2+ score and focal gene amplification was negative in the PDX suggesting clonal outgrowth [12].

The PDXs showed 4 distinct architectural patterns reminiscent of human breast disease: flat (F), lobular (LOB), *in situ* (IS), and invasive (INV) in pure form or mixed (Figure 2). The **F** pattern (Figure 2A-E), reminiscent of columnar cell changes and flat epithelial atypia, early premalignant alteration [11,12] was characterized by a monolayer lining of variably dilated mouse ducts (Figure 2A) with large columnar tumor cells showing mild nuclear pleomorphism and abundant eosinophilic cytoplasm forming apical “snouts” (Figure 2B). It was associated with foci of intraductal proliferation of low nuclear grade tumor cells in a cribriform pattern similar to atypical ductal hyperplasia [11] (Figure 2E). Strong and diffuse ER (Figure 2D) and PgR expression (Figure 2E) were observed in the F pattern, interpreted as an early step of “colonization” of the murine ducts by human tumor cells.

**Figure 2:**
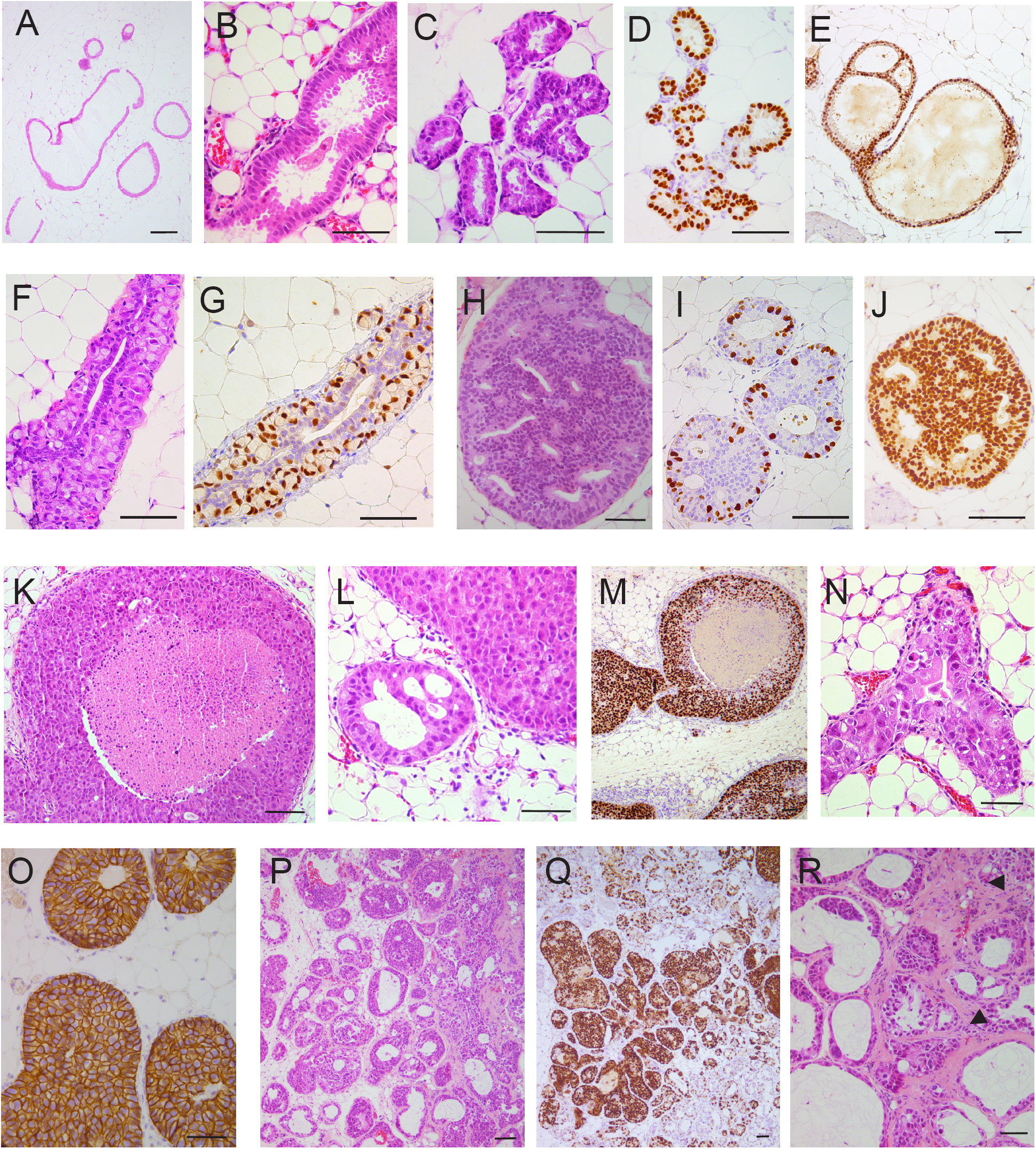
ID-PDXs exhibit four morphological patterns by H&E and IHC (A-E) Flat (F) pattern. (A-C) Human tumor cells cover the wall of variably dilated ducts forming a monolayer of large cylindrical cells with small nuclei, eosinophilic cytoplasm, and apical cytoplasmic “snouts” reminiscent of columnar cell changes and flat epithelial atypia in the human breast. D, E) Estrogen (ER) (D) and PgR (E) are strongly and diffusely expressed by human cells. E) Focally, transition between the F pattern and cellular bridges similar to human atypical ductal hyperplasia is seen. **(F-G) lobular (LOB) pattern.** (F) The mouse ductal epithelium is replaced by human cells harboring 2 associated phenotypes: i) large cells with a “signet ring cell” appearance, a large cytoplasmic vacuole displaces an enlarged nucleus toward the periphery of the cell, grow within the ductal wall in a “pagetoid” manner; ii) smaller cohesive cubo-cylindrical cells lining the ductal lumen. (G) The ER is expressed in most large cells and in a few cylindrical cells. **(H-M) In Situ (IS) pattern**. Human tumor cells fill the ductal lumen. (H-J) Some IS PDXs exhibit a cribriform architectural pattern, mild nuclear pleomorphism, low proliferation index (Ki67) (I), and diffuse and strong expression of the ER (J). (K-M) One PDX exhibits a solid growth with comedonecrosis, marked nuclear pleomorphism, high proliferation index (Ki67) (L), and HER2 overexpression (M). **(N-P) Invasive pattern (INV**): Tumor cells diffusely expressing the ER fill the dilated mouse ductal tree (N, O), and grow outside the ducts in small clusters surrounded by collagen (P, arrows). Scale bars: 100 μm.

The **LOB** pattern was characterized by tumor cell growth within the ductal wall, like pagetoid spread of lobular carcinoma *in situ* (LCIS) (Figure 2F, G) [11,13]. It was associated with intracellular clear, mucin-like vacuoles bestowing a signet-ring cell-like appearance on the tumor cells (Figure 2F). Interestingly, 4/5 lobular PDXs corresponded to this pattern, no typical LCIS was observed in the 5 lobular-PDXs, instead 4 of them exhibited the F pattern and 4 the LOB pattern. In the human breast also, columnar cell changes and lobular neoplasia are frequently associated in precursor lesions of “low grade” BC of both ductal and lobular subtypes [14,15]. The only case of pleomorphic lobular carcinoma showed LOB and F patterns (Fig. 1B) with a 20% proliferative fraction.

The **IS** pattern showed varying degrees of filling of the ductal lumen by tumor cells (Figure 2H-M) and resembled ductal carcinoma *in situ* (DCIS). It encompassed variable combinations of different degrees of nuclear pleomorphism and architectural patterns, like cribriform (Figure 2H-J) or solid with or without *comedo*necrosis (Figure 2K-M) [11]. An ER+ HER2+ BC-derived PDX showed large tumor extent, high nuclear pleomorphism, *comedo* architecture, high proliferation rate (Figure 2L), and HER2 overexpression (Figure 2M).

Although the tumor cells were derived from invasive primary tumors, only in 1/3 of the PDXs was of **INV** pattern with tumor cells detected outside the host ducts, either isolated or in small clusters (Figure 2N-P). Thus, intraductally xenografted tumor cells replicate earlier stages than the original PT. Notwithstanding the early developmental stages, micrometastases were present in 87.5% of the mice and in all the 16 analyzed PDXs (Table S1).

Next, we assessed how the biological features of the PDXs related to the prognostic characteristics of their clinical counterparts. According to a 20% Ki67 IHC cut-off based on clinical practice [16], the PT series comprised 7 *luminal A-like* (Ki67 <20%) and 14 *luminal B-like* (Ki67 ≥20%) cases; 4 of the latter were HER2+. In *luminal A-like* cases, the F pattern was observed in 6/7 PDXs, pure in 3 cases, mixed with IS in 1, and with LOB pattern in 2 cases. No INV pattern was found and the IS pattern only in one case (Table 1, Figure 3A). In *luminal B-like* cases, all NST, the F pattern was observed in 9/14 and only 2 were pure (Figure 3A). The INV pattern was detected in 6/14, IS and INV patterns were combined in 4/14. Thus, there was a trend for the F pattern to be present in PDXs obtained from *luminal A-like* BC. The IS and INV patterns on the contrary were observed, with or without associated F pattern, in PDXs obtained from the more aggressive *luminal B-like* subtype. Metastatic burden was significantly higher in mice engrafted with *luminal B-like* than *luminal A-like* PTs (Figure 3B).

**Figure 3:**
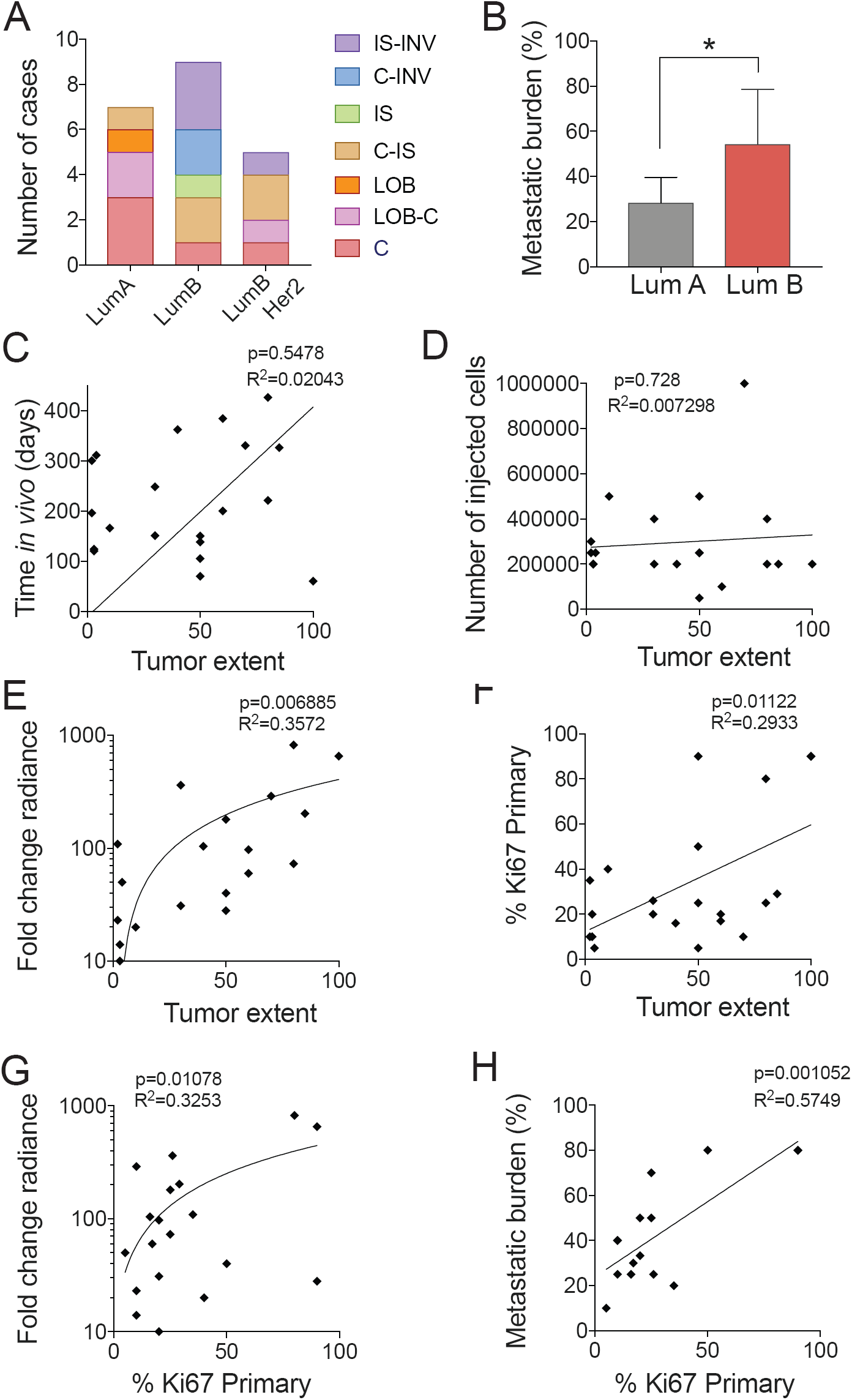
Characterization of *in vivo* growth. (A) Distribution of histological patterns within each subtype. (B) Metastatic burden expressed as percentage of positive organs in each recipient in Lum A versus Lum B derived PDXs. (C-F) Graph indicating Pearson correlation of tumor extent against (C) time of PDX growth *in vivo*, expressed in days after injection, (D) initial number of cells injected per gland, (E) *in vivo* cell growth expressed as fold change radiance relative to the first day of measurement (log10) and (F) Ki67 index of the primary tumor. (G,H) Pearson correlation of primary tumor Ki67% with (G) *in vivo* monitored growth and with (H) metastatic burden the percentage of organs affected by metastases within each case as determined by bioluminescence.

Tumor extent did not correlate with the time the grafted cells spent in the hosts (Figure 3C) nor with the number of cells injected (Figure 3D) but with *in vivo* growth rates (Figure 3E) and Ki67 index in PT (Figure 3F). The proliferative indices in PTs also correlated with *in vivo* growth rates (Figure 3G) and metastatic load (Figure 3H). Thus, tumor extent relates to PT biology rather than to engrafment modalities; growth rates and metastatic burden reflect Ki67 index and hence patient prognosis.

## Discussion

ID-PDXs of ER+ BCs exhibit a phenotypic diversity, which reflects the entire spectrum of human disease progression from precursor lesions to *in situ* disease and invasive cancers [11,13,17]. Our findings indicate that critical histopathologic features are intrinsic to tumor cells and not determined by the stroma or systemic factors, which differ between human and mice. The observation that two-thirds of the ID-PDXs derived from invasive cancers represent earlier lesions, like *in situ* carcinoma or flat epithelial atypia suggest that engrafted tumor cells re-start their growth intraductally, recapitulating different steps with, for some of them, a prolonged stage of non-invasive growth as they likely did in the patient’s breast years before diagnosis. This phenomenon is important to consider when using the model in personalized medicine [9].

While the typical F, LOB, IS patterns are readily identified, some morphological aspects of ER+ ID-PDX are more difficult to classify. This concerns dispersed foci of human cells lining the wall of small mouse ducts, with neither apical snouts nor cystic duct dilation. We classified these as F but speculate that they will develop into a different pattern.

While the majority (15/21) of PDXs have some F pattern, this pattern tends to remain pure in PDXs from luminal A PTs whereas IS and INV patterns are detected when luminal B cells were engrafted. Furthermore, the proliferative indices of the PT is maintained in the PDXs. Thus, the heterogeneity of luminal tumors is preserved in this model. This is especially important regarding the clinical problem of overtreatment; the model may help identify tumors with low aggressiveness and/or a high level of responsiveness to endocrine therapy.

## Figures legends

**Figure S1.**
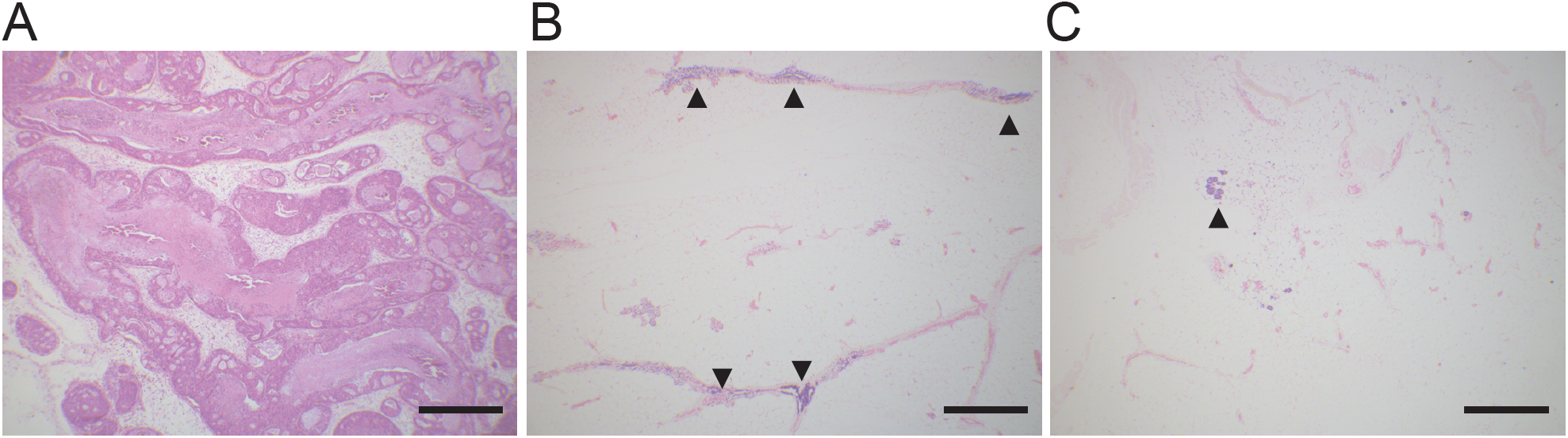
Different tumor extents in PDXs. (A) High tumor extent: Intraductally injected human tumor cells occupy over 95% of the mouse mammary ductal tree (H&E, x 4). (B) Moderate tumor extent: human tumor cells occupy between 10% and 50% of the mouse ductal tree (Alu staining, x 4). (C) Low tumor extent: human tumor cells form small foci occupying less than 5% of the mouse mammary ductal tree (ALU staining,). Scale bars: 800 μm.

**Figure S2.**
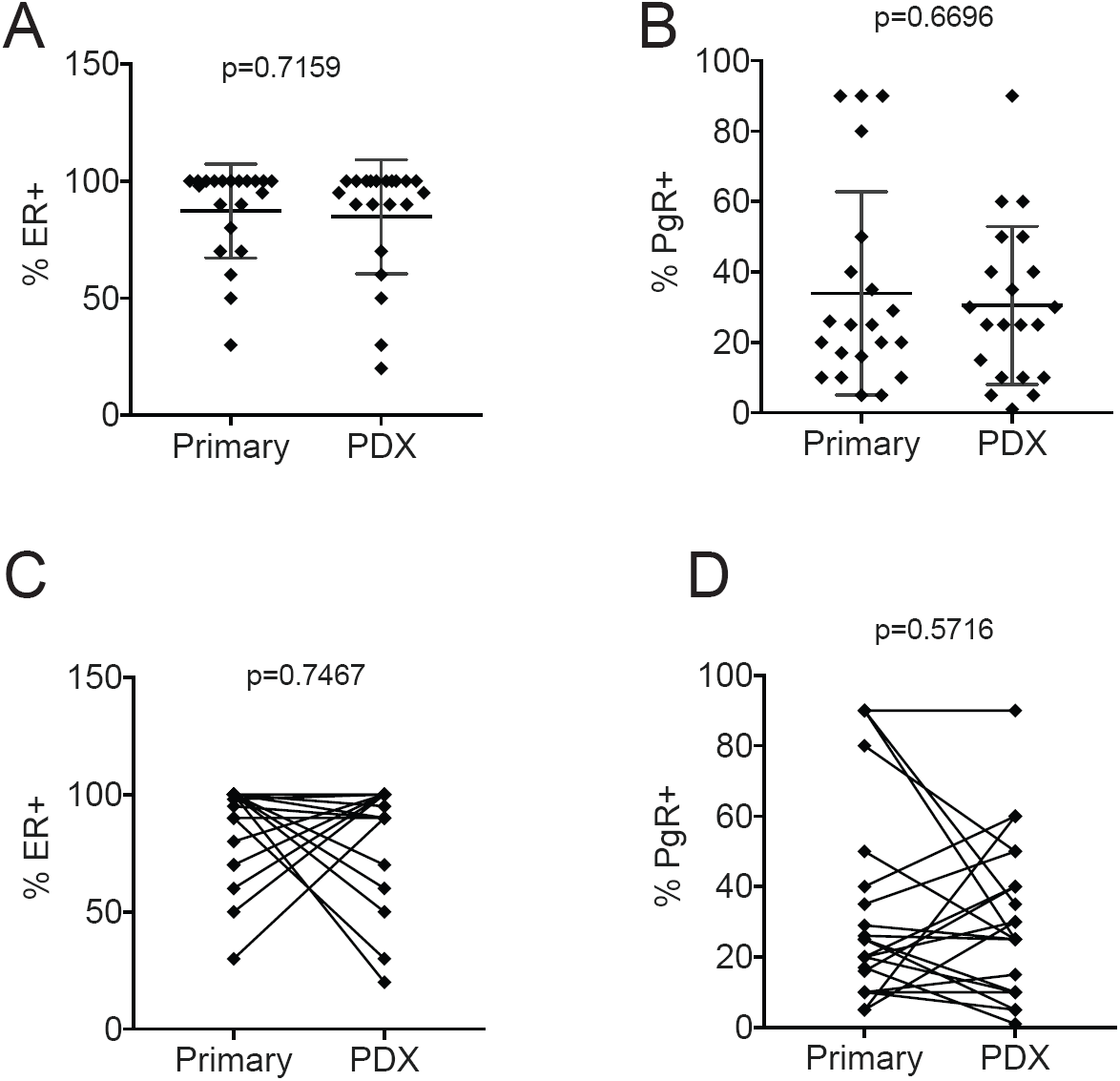
Marker expression. (A, C) Overall distribution of ER and PgR expression between PT and PDX, (B, D) Matched ER and PgR expression percentages of primary tumor and of the intraductally injected cells.

**Table S1 Characterization of primary tumors (PTs) and PDXs.** Information on patient, primary tumor, TNM stage, and histological types according to the 7^th^ edition of the TNM classification and WHO Classification of breast tumors [11]. Abbreviations : pos : positive; het : heterogeneous; w/m/s: weak/moderate/strong intensity of IHC. ND: not determined. NA: not applicable. AVG FC average fold change in radiance.

## Acknowledgements

We thank the Pathology Institute of the University Hospital of Lausanne (CHUV, Centre Hospitalier Vaudois) for its contribution to collecting and characterizing the primary tumors.

## Author contributions

AT, AS, KZ, and RLS selected patients for participation in the study, obtained informed consent, and participated in planification of tumor collection with LB. VS, AA, GS performed experiments and analyzed data. MF took tumor samples, MF and GS analysed PDXs sections, MF and CB wrote the manuscript. All authors were involved in writing the paper and had final approval of the submitted and published versions. The work was supported by the Swiss Cancer Ligue KFS-3701-08-2015 and by Biltema and ISREC Foundation.

